# Pancreatic Beta Cell Autophagy is Impaired in Type 1 Diabetes

**DOI:** 10.1101/2020.09.10.291443

**Authors:** Charanya Muralidharan, Abass M. Conteh, Michelle R. Marasco, Justin J. Crowder, Jeroen Kuipers, Pascal de Boer, Ben N.G. Giepmans, Amelia K. Linnemann

**Affiliations:** Departments of Biochemistry and Molecular Biology, Indianapolis, IN; Pediatrics, Indiana University School of Medicine, Indianapolis, IN; Department of Biomedical Sciences of Cells and Systems, University Medical Center Groningen, University of Groningen, Groningen, The Netherlands; Center for Diabetes and Metabolic Diseases, Indiana University School of Medicine, Indianapolis, IN

**Author notes:** Corresponding Author: Amelia K. Linnemann, Indiana University School of Medicine, 635 Barnhill Drive, Indianapolis, IN 46202.

**Keywords:** Type 1 diabetes, autoantibody positive, autophagy, crinophagy, autophagosome, lysosome

## Abstract

**Aims/hypothesis:** Pancreatic beta cells are highly metabolic secretory cells that are subjected to exogenous damaging factors such as proinflammatory cytokines or excess glucose that can cause accumulation of damage-inducing reactive oxygen species (ROS) during the pathogenesis of diabetes. We and others have shown that beta cell autophagy can reduce ROS to protect against apoptosis both *in vitro* and *in vivo.* While impaired islet autophagy has been demonstrated in human type 2 diabetes, it is unknown if islet autophagy is perturbed in the pathogenesis of type 1 diabetes. We hypothesized that beta cell autophagy is dysfunctional in type 1 diabetes, and that there is a progressive loss during early diabetes development.

**Methods:** Mouse pancreata were collected from chloroquine injected and non-injected NOR, nondiabetic NOD, and diabetic NOD mice. Age and BMI-matched pancreas tissue sections from human organ donors (n=34) were obtained from the Network for Pancreatic Organ Donors with Diabetes (nPOD). To assess autophagic flux, we injected the mice with chloroquine to inhibit the final stages of autophagy. We analyzed tissues for markers of autophagy via immunofluorescence analysis. Tissue sections were stained with antibodies against proinsulin or insulin (beta cell markers), LC3A/B (autophagosome marker), Lamp1 (lysosome marker), and p62 (autophagy adaptor protein and marker for autophagic flux). Images were collected on a scanning laser confocal microscope then analyzed with CellProfiler and ImageJ. Secondary lysosomes and telolysosomes (formerly called lipofuscin bodies, residual bodies or tertiary lysosomes) were analyzed in electron micrographs of pancreatic tissue sections from human organ donors (nPOD; n=12) deposited in www.nanotomy.org/OA/nPOD. Energy Dispersive X-ray (EDX) analysis was also performed on these tissues to analyze distribution of elements such as nitrogen, phosphorus, and osmium in secondary lysosomes and telolysosomes of nondiabetic and autoantibody positive donor tissues (n=5).

**Results:** We observed increased autophagosome numbers in islets of diabetic NOD mice (p=0.0077) and increased p62 in islets of both nondiabetic and diabetic NOD mice (p<0.0001 in both cases) when compared to NOR mice. There was also a significant reduction in autophagosome:lysosome colocalization in islets of diabetic NOD mice compared to both nondiabetic NOD mice (p=0.0004) and NOR mice (p=0.0003). Chloroquine infusions elicited accumulation of autophagosomes in the islets of NOR (p=0.0029) and nondiabetic NOD mice (p<0.0001), but not in the islets of diabetic NOD mice. Chloroquine also stimulated an accumulation of the autophagy adaptor protein p62 in the islets of NOR mice (p<0.001), however this was not observed in NOD mice (regardless of diabetes status). In the human pancreata, we observed significantly reduced autophagosome:lysosome colocalization (p=0.0002) in the residual beta cells of donors with type 1 diabetes compared to nondiabetic controls. We also observed reduced colocalization of proinsulin with lysosomes in the residual beta cells of donors with type 1 diabetes compared to both nondiabetic (p<0.0001) and autoantibody positive donors (p<0.0001). Electron microscopy based analysis of human pancreas sections also revealed accumulation of telolysosomes in beta cells of autoantibody positive donors (p=0.0084), the majority of which had an nitrogen dense ring outside a phospholipid core.

**Conclusions/interpretation:** Collectively, we provide evidence of impairment in the final degradation stages of islet macroautophagy and crinophagy in human type 1 diabetes. We also document an accumulation of telolysosomes with nitrogen accumulation at their periphery in the beta cells of autoantibody positive donors. This demonstrates clear differences in the lysosome contents of autoantibody positive donors that may be associated with lysosome dysfunction prior to clinical hyperglycemia. We observe similar impairments in macroautophagy in the diabetic NOD mouse, a model of type 1 diabetes, suggesting that this mouse model can be appropriately used to study the pathogenesis of autophagy/crinophagy loss and how it relates to disease initiation and progression. Considering these data in the context of what is known regarding the cell-protective effects of islet autophagy, we suggest targeting beta cell autophagy pathways as an approach to reduce apoptosis in individuals at risk for type 1 diabetes development.

**Research in context:** *What is already known about this subject?*

- Autophagy confers a cytoprotective role in the beta cell.
- Autophagy is reduced in type 2 diabetes.
- Autophagy in the context of type 1 diabetes is unexplored.

*What is the key question?*

- Is autophagy reduced during the pathogenesis of human type 1 diabetes?

*What are the new findings?*

- We provide evidence of reduced autophagy and crinophagy in human type 1 diabetes.
- These data are supported by observations of reduced autophagy in a mouse model of autoimmune diabetes.

*How might this impact on clinical practice in the foreseeable future?*

- This study provides evidence that autophagy is impaired in human type 1 diabetes. Prior studies have shown that loss of autophagy in the islet is associated with increased beta cell apoptosis, therefore we propose that therapeutic targeting of autophagy pathways may reduce beta cell death in type 1 diabetes development.

## Introduction

Beta cell ROS accumulation has been implicated as a triggering event during the development of type 1 diabetes [1]. ROS can be generated from multiple sources within a cell, such as mitochondrial oxidative phosphorylation, metabolism of long chain fatty acids in the peroxisome, and enzyme catalysis [2]. Although low levels of ROS can promote important signaling events in the beta cell such as insulin secretion [3] and proliferation [4], excess ROS can damage cellular proteins and organelles and overwhelm the endogenous mechanisms that maintain homeostasis [5].

Autophagy is an endogenous mechanism to reduce ROS and promote beta cell survival [6]. Known types of autophagy, including macroautophagy, chaperone mediated autophagy, microautophagy, and crinophagy function as important housekeeping catabolic processes to facilitate recycling of excess or damaged cellular components and promote cellular homeostasis. Macroautophagy (herein referred to as autophagy) is a dynamic process that involves a cascade of regulated events leading to engulfment of damaged proteins or organelles into the microtubule associated protein 1 light chain 3 (MAP1LC3, or LC3)-containing double membraned autophagosome. Mature autophagosomes fuse with acidic lysosomes for cargo degradation and recycling. Crinophagy is a specialized form of “-phagic” degradation that occurs in secretory cells, where the secretory granules fuse directly with the lysosome [7]. Although the mechanism of crinophagy is not well-understood, this process plays a key role in the regulation of insulin granules [8].

The critical importance of autophagy for beta cell homeostasis and survival in the context of type 2 diabetes has been well-documented [9]. Although there is no literature demonstrating impaired autophagy in the context of human type 1 diabetes, dysfunctional autophagy has been implicated in the pathogenesis of several autoimmune disorders [10], suggesting a potential role in type 1 diabetes pathogenesis. Indeed, a role for the autophagic pathway in type 1 diabetes-associated autoimmunity has been hypothesized [11]. We thus aimed to assess if there is a decline in islet autophagy associated with type 1 diabetes.

## Methods

### Mice

Nonobese diabetes Resistant (NOR) and Nonobese Diabetic (NOD) mice were purchased from Jackson Labs at ~7 weeks of age. Mice were housed in a temperature-controlled facility with a 12-h light/12-h dark cycle and were given free access to food and water. All experiments were approved by the Indiana University School of Medicine Institutional Animal Care and Use Committee. Blood glucose for NOR mice and NOD mice was monitored bi-weekly with an AlphaTrak2 glucometer, and NOD mice were characterized as diabetic after 2 consecutive days of blood glucose readings >250 mg/dL. Mice were euthanized by cervical dislocation and pancreata were collected. Harvested tissues were fixed in 3.7% paraformaldehyde (v/v) for 4-5 hours at room temperature with gentle agitation and then transferred to 70% ethanol. Pancreata were then paraffin embedded and sectioned in the Histology Core of the Indiana Center for Musculoskeletal Health, Indiana University School of Medicine. For all experiments, female mice aged 12-26 weeks were used.

### Chloroquine injections

To analyze dynamic autophagic flux, a subset of NOR mice (12 weeks; n=5), non-diabetic NOD mice (14 weeks; n=5), and diabetic NOD mice (14-26 weeks; n=3) were intra-peritoneally injected with 50mg/kg of Chloroquine diphosphate (Tocris #4109). Two hours post injection, pancreata were collected, paraffin embedded and sectioned. For non-injected controls, a subset of NOR mice (n=5), nondiabetic NOD (n=6), and diabetic NOD (n=7) were used.

### Human Organ Donor characteristics

We obtained deidentified pancreatic tissue sections from 32 human organ donors through the JDRF Network for Pancreatic Organ Donors with Diabetes (nPOD; **Table 1**). Samples included sections from 12 non-diabetic control donors (6 males, 6 females), 12 autoantibody positive donors (6 males, 6 females) and 10 donors with type 1 diabetes (5 males, 5 females) that had residual insulin positive islets. Age- (mean±SEM= 26.15±1.081) and Body Mass Index- (BMI; mean±SEM= 26.55±0.789) matched donor samples were used. Breakdown of age and BMI in each group is shown in **ESM Fig.1**. The duration of diabetes ranged from <1 year to 32.5 years.

**Table 1.** nPOD donor characteristics

| c | Donor ID | Age (years) | Sex | BMI | C-peptide | T1D duration (years) | Autoantibody |
| --- | --- | --- | --- | --- | --- | --- | --- |
| Nondiabetic | 6015 | 39 | F | 32.2 | 1.99 | - | - |
| Nondiabetic | 6160 | 22.1 | M | 23.9 | 0.4 | - | - |
| Nondiabetic | 6162 | 22.7 | M | 28.9 | 7.61 | - | - |
| Nondiabetic | 6178 | 24.5 | F | 27.5 | 4.55 | - | - |
| Nondiabetic | 6235 | 30 | M | 25.4 | 8.1 | - | - |
| Nondiabetic | 6335 | 18.8 | M | 23.6 | 8.85 | - | - |
| Nondiabetic | 6339 | 23.3 | M | 25 | 10.56 | - | - |
| Nondiabetic | 6373 | 15.7 | M | 25 | 13.12 | - | - |
| Nondiabetic | 6034 | 32 | F | 25.2 | 3.15 | - | - |
| Nondiabetic | 6234 | 20 | F | 25.6 | 6.89 | - | - |
| Nondiabetic | 6331 | 27.1 | F | 24 | 3.01 | - | - |
| Nondiabetic | 6333 | 27.1 | F | 24.9 | 9.37 | - | - |
| Nondiabetic | 6226 | 38 | F | 27.2 | 3.87 | - | - |
| Nondiabetic | 6227 | 17 | F | 26.4 | 2.75 | - | - |
| Nondiabetic | 6229 | 31 | F | 26.9 | 6.23 | - | - |
| Nondiabetic | 6230 | 16 | M | 19.4 | 5.22 | - | - |
| Nondiabetic | 6232 | 14 | F | 20.83 | 19.5 | - | - |
| Aab+ | 6147 | 23.8 | F | 32.9 | 3.19 | - | GADA |
| Aab+ | 6167 | 37 | M | 26.3 | 5.43 | - | IA2A, ZnT8A |
| Aab+ | 6151 | 30 | M | 24.2 | 5.49 | - | GADA |
| Aab+ | 6170 | 34.5 | F | 36.9 | 4.29 | - | GADA |
| Aab+ | 6181 | 31.9 | M | 21.9 | 0.06 | - | GADA |
| Aab+ | 6301 | 26 | M | 32.1 | 3.92 | - | GADA |
| Aab+ | 6310 | 28 | F | 22.4 | 10.54 | - | GADA |
| Aab+ | 6314 | 21 | M | 23.8 | 1.49 | - | GADA |
| Aab+ | 6397 | 21.16 | F | 29.6 | 12.77 | - | GADA |
| Aab+ | 6400 | 25.15 | M | 22.2 | 4.17 | - | GADA |
| Aab+ | 6450 | 22 | F | 24.4 | 5.47 | - | GADA, ZnT8A |
| Aab+ | 6483 | 30.46 | F | 20.8 | 2.21 | - | GADA |
| Aab+ | 6388 | 25.2 | F | 26 | 1.38 | - | GADA, mIAA |
| Aab+ | 6303 | 22 | M | 31.9 | 3.03 | - | GADA |
| Aab+ | 6197 | 22 | M | 28.2 | 17.48 | - | GADA, IA2A |
| Aab+ | 6156 | 40 | M | 19.8 | 13.34 | - | GADA |
| T1D | 6046 | 18.8 | F | 25.2 | 0.05 | 8 | IA2A, ZnT8A |
| T1D | 6302 | 38.5 | M | 20.5 | 0.17 | 32.5 | - |
| T1D | 6306 | 19 | M | 24.5 | 0.02 | 5 | mIAA |
| T1D | 6325 | 20 | F | 31.2 | 0.14 | 6 | GADA, IA2A, mIAA |
| T1D | 6328 | 39 | M | 24 | 0.02 | 20 | GADA, mIAA |
| T1D | 6367 | 24 | M | 25.7 | 0.39 | 2 | - |
| T1D | 6405 | 29.1 | F | 42.5 | 1.84 | 0.6 | GADA, IA2A, ZnT8A |
| T1D | 6414 | 23.1 | M | 28.4 | 0.16 | 0.43 | GADA, mIAA, ZnT8A |
| T1D | 6435 | 24.75 | F | 26.9 | 0.09 | 14.75 | - |
| T1D | 6477 | 19.87 | F | 25.3 | 0.18 | 8 | GADA, IA2A, mIAA |

### Immunofluorescence analysis

Tissue sections were dewaxed in xylene and hydrated in serial dilutions of ethanol (2x; 100%, 90% and 70%) followed by water. Heat and citrate-based antigen retrieval was performed using Vector Laboratories antigen unmasking solution (H-3300) for 20 minutes (in 5-minute bursts) and then the slides were allowed to cool down to room temperature for 1 hour. Slides were blocked with Dako blocking buffer (Agilent Technologies X0909) prior to incubation with antibodies in Dako antibody diluent (Agilent technologies S3022). Antibodies and the corresponding dilutions used are listed as follows: Developmental Studies Hybridoma Bank (DSHB) mouse anti-proinsulin (GS-9A8; 1:50), rat anti-Lamp1 (DSHB 1D4B; 1:50), rabbit anti-LC3A/B (Cell Signaling Technology 12741; 1:100), rabbit anti-p62 (Abcam ab91526; 1:200), and guinea pig anti-insulin (BioRad 5330-0104G; 1:500). Highly cross adsorbed fluorescently conjugated secondary antibodies were used. Images were collected on either a Zeiss LSM 700 confocal microscope using a 63X/1.4 NA oil objective, or a Zeiss LSM 800 confocal microscope equipped with Airyscan using a 63X/1.4 NA oil objective.

### Cell profiler analysis flow of logic

Fluorescence intensity measurements, puncta counts, and colocalization analyses were automated using Cell Profiler versions 3.1.5/8/9 [12]. Background from each image was first subtracted by removing the lower quartile intensity across each channel. Regions of interest (ROI) for human pancreatic tissues were defined by proinsulin positive area and ROIs for mouse pancreatic tissues were defined by manually identifying the islet area. Puncta for LC3, Lamp1, p62 and proinsulin were identified by discarding objects outside a pixel diameter range, after applying median filtering (**ESM Fig. 2**). For human islets, the puncta count was normalized to number of nuclei in the proinsulin positive area. For mouse islets, the puncta count was normalized to the area of the islet ROI. For colocalization analysis, parent and child objects were defined. For example, for LC3 puncta colocalization analysis with Lamp1 puncta, Lamp1 and LC3 were defined as parent and child objects, respectively, and the percentage of LC3 puncta cocompartmentalizing with Lamp1 puncta was determined by following the cell profiler tutorial for object overlap based colocalization [13].

### Human Electron Microscopy data analysis

Electron microscopy (EM) images from control and autoantibody positive nPOD donor pancreatic tissue sections were analyzed from the Nanotomy repository ([14]; **Table 1**). At least 48 beta cells were analyzed per group. Structures that morphologically resemble lipofuscin bodies, as described by Cnop *et al* [15], were quantified within beta cells. Lipofuscin bodies are also named residual bodies, tertiary lysosomes and more recently telolysosomes. Elemental analysis of nitrogen, osmium, and phosphorus for telolysosomes and secondary lysosomes in nondiabetic (n=2) and autoantibody positive (n=3) donor sections was performed using EM-Energy dispersive X-ray analysis (EDX) as described previously [16] on cells which could be identified with certainty as beta cells by the presence of insulin granules.

### Statistical analysis

At least 47 proinsulin positive islets were analyzed from the pancreatic tail region for each donor group and at least 18 islets were analyzed for each mouse group, with representative images shown (**Figures 1-4**). For EM image analysis, at least 48 insulin positive beta cells were analyzed from the pancreatic head or body region for each donor group, with representative images shown in **Figure 5**. Data analyzed using Cell Profiler were compared between groups by one-way ANOVA with multiple comparisons (Tukey’s post hoc test), two-way ANOVA with multiple comparisons (Sidak’s post hoc test), or unpaired t-test, as deemed appropriate, using Graphpad Prism v8.0 and are represented as mean±SEM. P-values < 0.05 were considered statistically significant.

**Figure 1.**
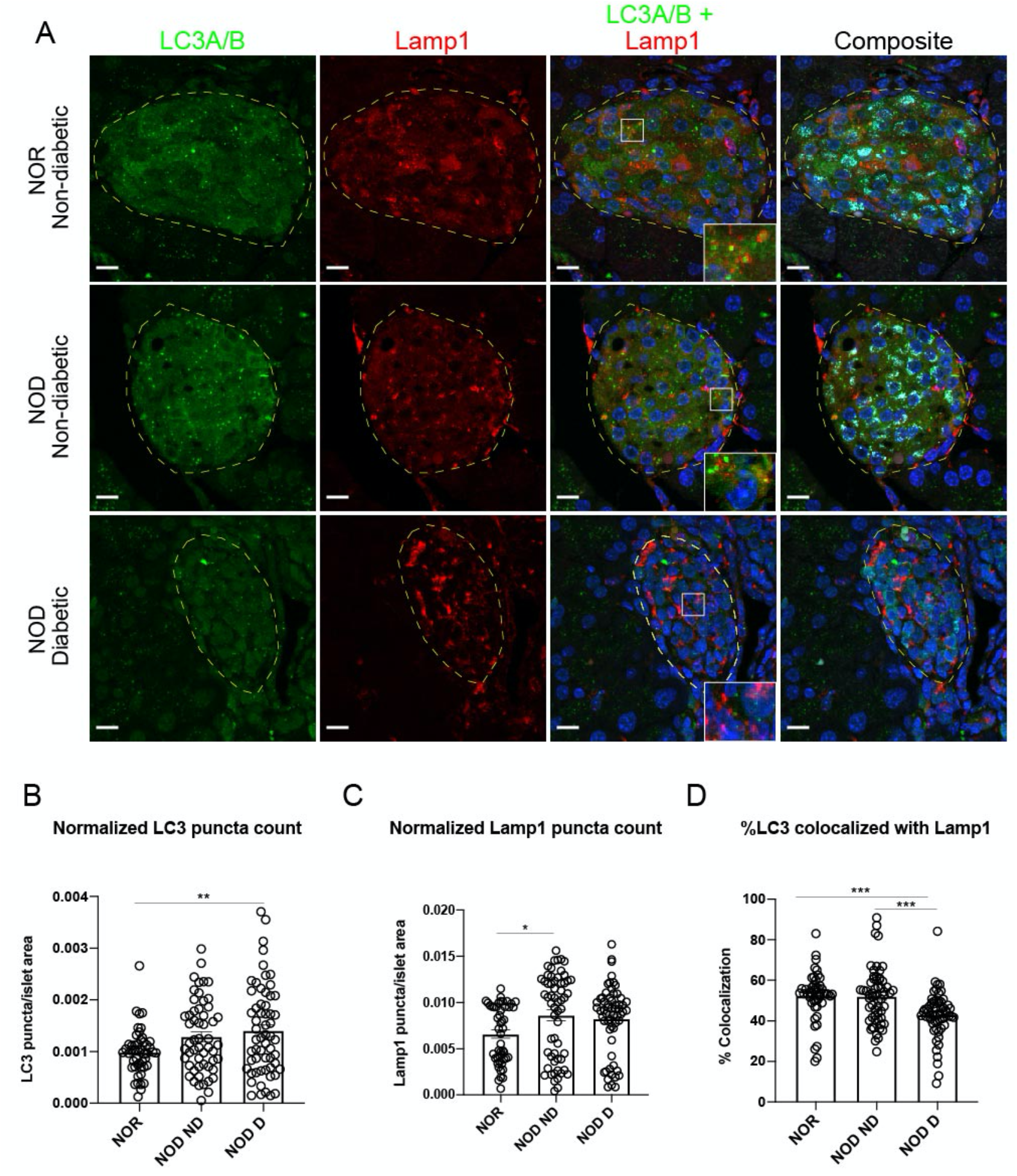
Reduced autophagy in the pancreatic islets of diabetic NOD mice. **(A)** Representative images showing immunofluorescent staining of islets in pancreas tissue sections from nondiabetic NOR, nondiabetic NOD and diabetic NOD mice. Autophagosomes (LC3A/B) are shown in green, lysosomes (Lamp1) in red, proinsulin in cyan, and nuclei (DAPI) in blue. Scale bars, 10μm. Insets show higher magnification of overlapping puncta. **(B)** Quantification of autophagosomes **(C)** Quantification of lysosomes **(D)** Quantification of colocalization of autophagosomes with lysosomes in islet. Each circle denotes an islet. *p<0.05; **p<0.01; ***p<0.001 (one-way ANOVA).

## Results

### Evidence of impaired autophagy in diabetic NOD mice

To determine if there is an impairment in autophagy in a mouse model of spontaneous autoimmune diabetes, we analyzed pancreata from diabetic NOD mice comparing them to both nondiabetic NOD mice and NOR mice, which do not develop insulitis or diabetes and are MHC-matched to NOD mice [17]. Representative immunofluorescent images are shown in **Figure 1A**. We observed a significant increase in LC3 puncta (autophagosome marker) in diabetic NOD mouse islets (p=0.0077), and a trend towards increased autophagosomes in nondiabetic NOD islets (p=0.0831) when compared to islets of NOR mice (**Fig. 1B**). This was accompanied by a significant increase in Lamp1 puncta (lysosome marker) in nondiabetic NOD mice (p=0.0186) and a trend towards an increase in the diabetic NOD mice (p=0.0670) compared to NOR mice (**Fig. 1C**).

The final step of autophagy is the degradation of autophagosomes and their cargo by lysosomal acid hydrolases. Therefore, we reasoned that if autophagy is impaired in the context of spontaneous autoimmune diabetes, this would be evident through altered autophagosome-lysosome colocalization. To assess defects in autolysosome formation or autophagosome-lysosome fusion, we therefore quantified the percentage of LC3 colocalized with Lamp1. We observed a significant reduction in the colocalization of LC3 with Lamp1 in diabetic NOD mouse islets when compared to either NOR or nondiabetic NOD mouse islets (p=0.0004 and p=0.0003, respectively; **Fig. 1D**), suggesting a defect in autophagosome-lysosome fusion during late stages of autophagy. Collectively, these data suggest an impairment in macroautophagy in the islets of diabetic NOD mice.

### Autophagic flux is impaired in diabetic NOD mice

Although we observed a reduction in the percentage of autophagosomes fusing with lysosomes in diabetic NOD mouse islets, these data do not necessarily capture the dynamic nature of the autophagic degradation process. Therefore, to assess autophagic flux and characterize basal autophagy status in NOD mice, we injected NOR mice and NOD mice (nondiabetic and diabetic) with chloroquine, a lysomotrophic drug that impairs the fusion of autophagosome and lysosomes [18] thus halting the degradation of formed autophagosomes. Our experimental approach is shown in **Figure 2A**, and representative immunofluorescent images of pancreata from chloroquine injected mice are shown in **Figure 2B**. To ensure that the chloroquine functioned as intended and to validate our colocalization analysis, we compared the islets of non-injected and chloroquine injected NOR and NOD (nondiabetic and diabetic) mice. We observed a statistically significant reduction in the colocalization of LC3 with Lamp1 in islets of chloroquine injected mice when compared to non-injected animals across all three groups (**Fig. 2C**), indicating that autophagosome:lysosome fusion was disrupted by the chloroquine, and suggesting that colocalization analysis is an appropriate readout for autolysosomes in immunofluorescence assays. We also observed a significant increase in islet LC3 puncta in chloroquine injected NOR (p=0.0029) and nondiabetic NOD mice (p<0.0001) compared to their respective non-injected controls (**Fig. 2D**), suggesting increased formation of autophagosomes in NOR and nondiabetic NOD mice. In contrast, there was no difference in islet LC3 puncta count in chloroquine injected diabetic NOD mice vs. non-injected control (**Fig. 2D**), further supporting our observation of defective autophagic flux in these mice. Additionally, we observed a significant increase in islet lysosome numbers in chloroquine injected NOR mice (p=0.0018; **Fig. 2E**). However, there was no difference in lysosomes of chloroquine injected NOD mice vs their corresponding non-injected controls, regardless of diabetes status (**Fig. 2E**).

**Figure 2.**
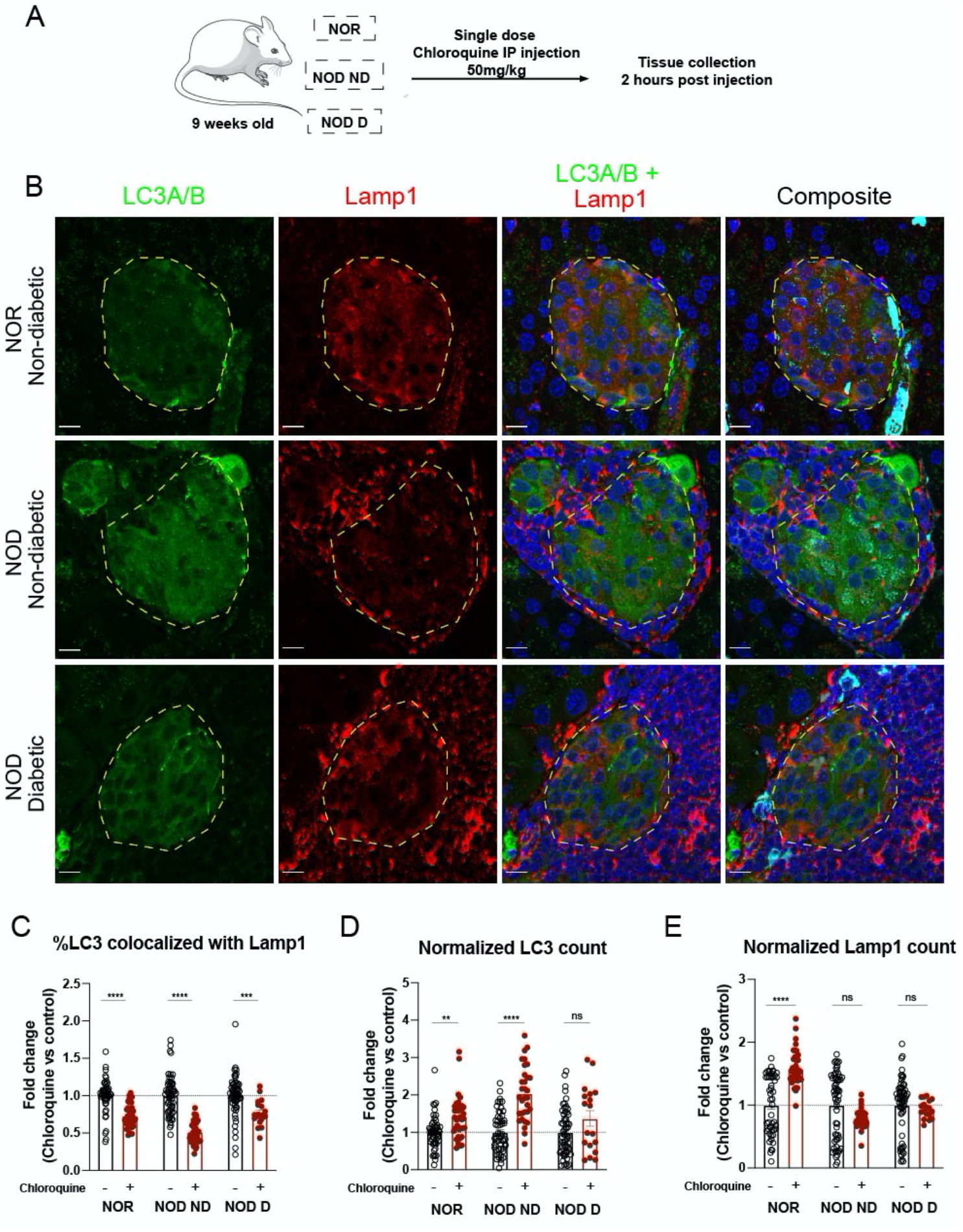
Impaired autophagic flux in the islets of diabetic NOD mice. **(A)** Schematic of autophagic flux assessment experiment. **(B)** Representative images showing immunofluorescent staining of islets of pancreas tissue sections from chloroquine injected nondiabetic NOR, and nondiabetic NOD. Autophagosomes (LC3A/B) are shown in green, lysosomes (Lamp1) in red, and nuclei (DAPI) in blue. Scale bars, 10μm. **(C)** Quantification of colocalization of autophagosomes with lysosomes in islet. **(D)** Quantification of autophagosomes. **(E)** Quantification of lysosomes. Each circle denotes an islet. *p<0.05; **p<0.01; ***p<0.001; ****p<0.0001 (two-way ANOVA with multiple comparisons).

Next, we analyzed p62 in islets of NOR and NOD mice (**Fig. 3A**). p62 is an autophagy adaptor protein that binds to both LC3 and to other proteins being targeted for degradation to aid in bringing them together [19], and is typically then degraded with the lysosomal contents during the autophagic degradation process [20]. Therefore, one would expect a reduction in p62 if autophagic flux is intact [21, 22]. At baseline, we observed an accumulation of p62 in islets of both nondiabetic and diabetic NOD mice (p<0.0001 in both cases) when compared to NOR mice (**Fig. 3B**). Chloroquine injection led to a significant increase in p62 levels in islets of NOR mice (p<0.0001) whereas there was no change elicited by chloroquine in NOD mice, regardless of diabetes status (**Fig 3C**). These data collectively suggest an impairment in autophagic flux in the islets of NOD mice that is more pronounced in the residual islets of diabetic NOD mice. Our analyses are also suggestive of impairment in the final stage of the autophagic degradation process in the context of spontaneous autoimmune diabetes in mice.

**Figure 3.**
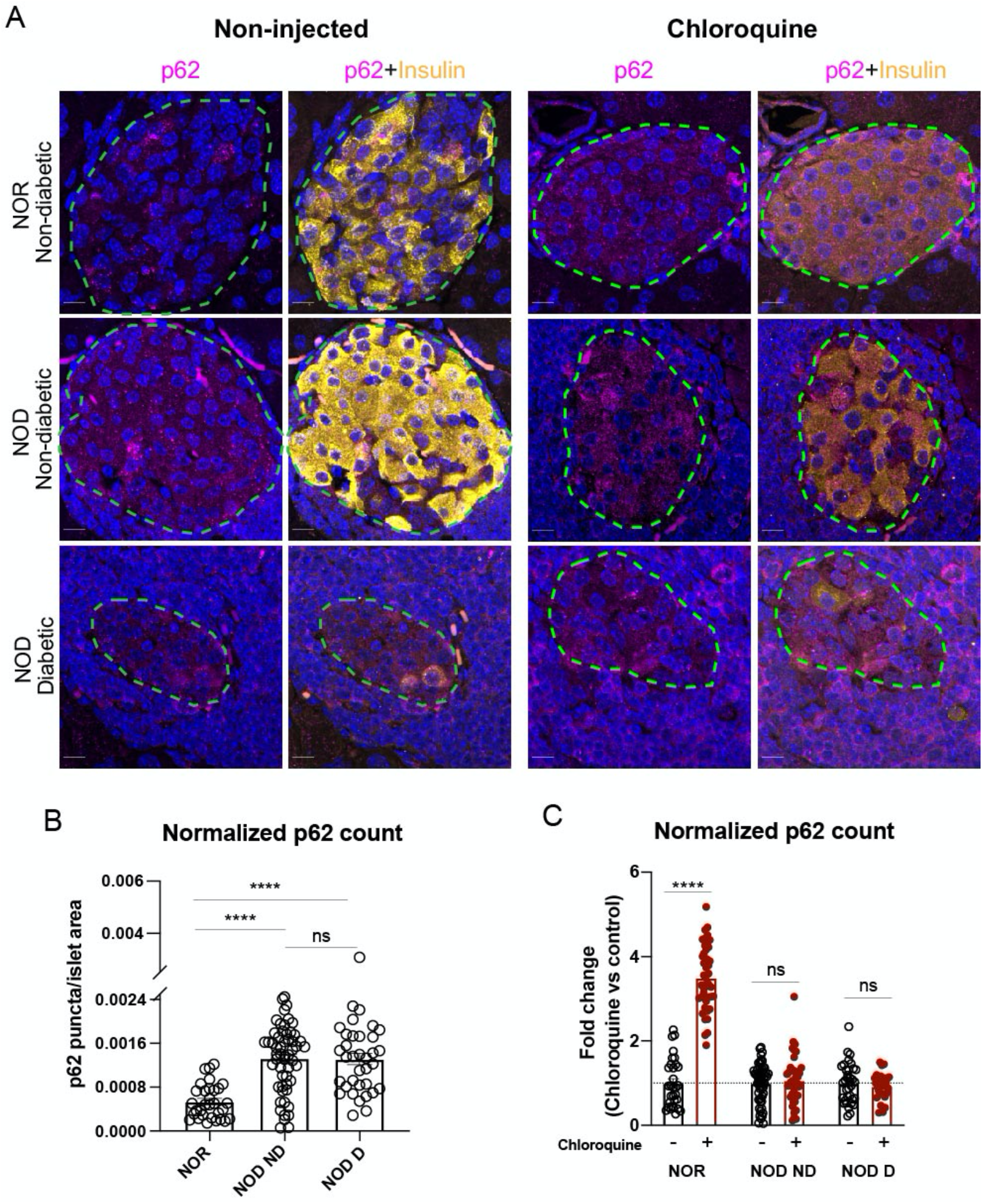
Impaired autophagic flux in nondiabetic and diabetic NOD mice islets. **(A)** Representative images showing immunofluorescent staining of islets of chloroquine injected NOR and NOD mice pancreatic tissue sections. p62 in magenta, Insulin in yellow and nuclei (DAPI) in blue. Scale bars, 10μm. **(B)** Quantification of p62 puncta in islet area in noninjected animals. **(C)** Fold change comparison of p62 puncta in islet area of chloroquine injected animals. Each circle denotes an islet. *p<0.05; **p<0.01; ***p<0.001; ****p<0.0001 (twoway ANOVA with multiple comparisons).

### Autophagosome numbers are reduced in beta cells of human type 1 diabetes donors

Impairment in autophagy has previously been demonstrated in the context of human type 2 diabetes [23, 24], but not in human type 1 diabetes. Therefore, we obtained human pancreata through nPOD. We first assessed the LC3A/B (autophagosome) puncta and Lamp1 (lysosome) puncta within the proinsulin-positive cells of nondiabetic control, autoantibody positive, or type 1 diabetic organ donor pancreata (**Figure 4A**). We observed a significant reduction of autophagosomes in beta cells of donors with type 1 diabetes when compared to autoantibody positive donors (p=0.0422; **Fig. 4B**). Interestingly, we also observed a trend towards increased autophagosome numbers in the beta cells of autoantibody positive donors when compared to beta cells of nondiabetic donors (p=0.0844; **Fig. 4B**). This data suggests possible accumulation of autophagosomes prior to the onset of clinical hyperglycemia. Similarly, lysosome numbers were significantly increased (p=0.0327) in the beta cells of donors with type 1 diabetes when compared to nondiabetic donors (**Fig. 4C**).

**Figure 4.**
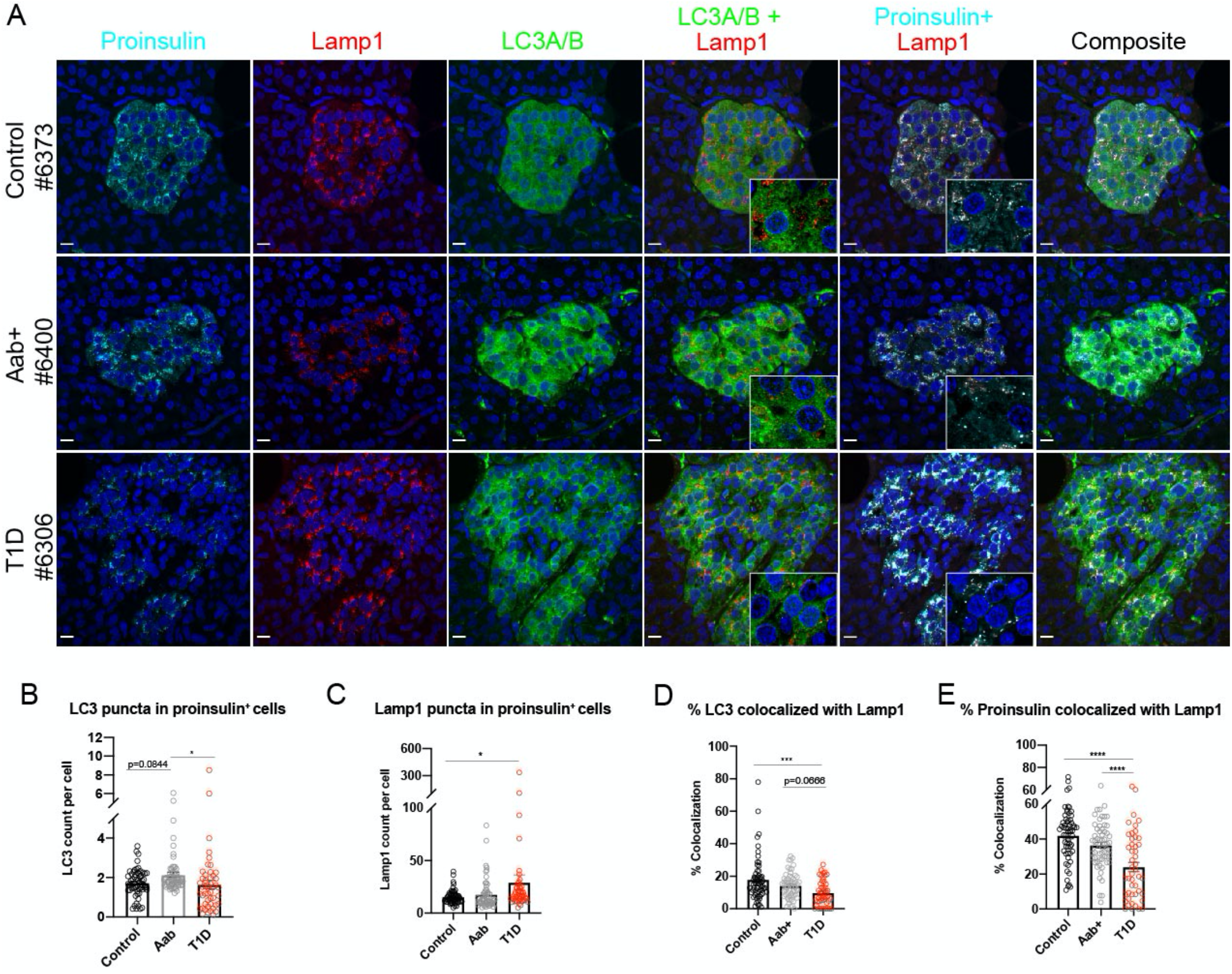
Reduced autophagy and crinophagy in the pancreatic beta cells of individuals with type 1 diabetes. **(A)** Representative images showing immunofluorescent staining of islets in proinsulin-positive cells (cyan) of pancreas tissue sections from nondiabetic, autoantibody positive (Aab+), and type 1 diabetic organ donors. Autophagosomes (LC3A/B) are shown in green, lysosomes (Lamp1) in red, and nuclei (DAPI) in blue. Scale bars, 10μm. Insets show higher magnification of overlapping puncta. **(B)** Quantification of lysosomes. **(C)** Quantification of autophagosomes. **(D)** Quantification of colocalization of autophagosomes with lysosomes in proinsulin-positive cells. **(E)** Quantification of colocalization of proinsulin with lysosomes in proinsulin-positive cells. Each circle denotes an islet. *p<0.05; **p<0.01; ***p<0.001; ****p<0.0001 (One-way ANOVA with multiple comparisons).

### Autophagy is reduced in human type 1 diabetes

We next quantified the colocalization of LC3 (autophagosome) puncta with Lamp1 (lysosome) puncta in the proinsulin-positive islet cells of nondiabetic, autoantibody positive, and type 1 diabetic human organ donors (**Fig. 4D**). We did not observe a statistically significant difference in autophagosome-lysosome colocalization between nondiabetic and autoantibody positive individuals. However, we observed a significant reduction in colocalization of autophagosomes with lysosomes in the residual beta cells of type 1 diabetic donors compared to nondiabetic donors (p=0.0002), and a trend towards decreased colocalization in the beta cells of type 1 diabetic donors vs autoantibody positive donors (p=0.0666; **Fig. 4D**). Additionally, there was no correlation between % LC3-lamp1 colocalization and disease duration or c-peptide levels (**ESM Fig. 3**). This demonstrates that autophagosome-lysosome fusion, and thus one of the final stages of autophagy, is impaired in the context of type 1 diabetes.

### Crinophagy is reduced in human type 1 diabetes

Crinophagy is another method of “-phagic” degradation by which the secretory granule directly fuses with the lysosomes. Crinophagy has been implicated in the presentation of altered peptides that recruit T-cells to the beta cell [25]. To determine if there is alteration of crinophagy in type 1 diabetes, we quantified the amount of proinsulin that colocalized with lysosomes (**Fig. 4E**). We observed a significant reduction in proinsulin-lysosome colocalization in islets of type 1 diabetic donors when compared to both nondiabetic (p<0.0001) and autoantibody positive individuals (p<0.0001). Similar to autophagy, we did not observe a correlation between crinophagy and disease duration or c-peptide levels (**ESM Fig. 3**). These data support the conclusion that crinophagy is also reduced in the context of type 1 diabetes.

### Telolysosomes are increased in autoantibody positive individuals

Autophagy and crinophagy are dependent on several factors, including the integrity of the lysosome acidification machinery to create an acidic environment that is capable of protein and organelle degradation. Telolysosomes (also known as lipofuscin bodies, residual bodies, or tertiary lysosomes) are lysosomes that contain undigestible biological garbage or cellular components such as highly oxidized, covalently crosslinked proteins, sugars or lipids [26]. Lipofuscin bodies have been shown to accumulate within post-mitotic cells that are long-lived and are positively correlated with aging in the beta cell [27]. Additionally, oxidative stress and inhibition of lysosomal enzymes have been associated with increased accumulation of lipofuscin bodies [28]. Therefore, we took advantage of electron microscopic images of nPOD pancreas tissue in the nanotomy repository [14], available from www.nanotomy.org/OA/nPOD. We identified telolysosomes (lipofuscin bodies) as described [15] (**Fig. 5A**) and quantified them in a series of samples from nondiabetic and autoantibody positive organ donors with some of the samples that were also analyzed by immunofluorescent analysis (**Fig. 5B**). We observed a significant increase in the number of telolysosomes in the beta cells of autoantibody positive individuals when compared to non-diabetic controls (p=0.0084). This suggests that the beta cells of autoantibody positive individuals are more prone to oxidative stress induced damage and therefore cell death, and suggests that autophagy may be impaired early in type 1 diabetes pathogenesis prior to the development of hyperglycemia.

**Figure 5.**
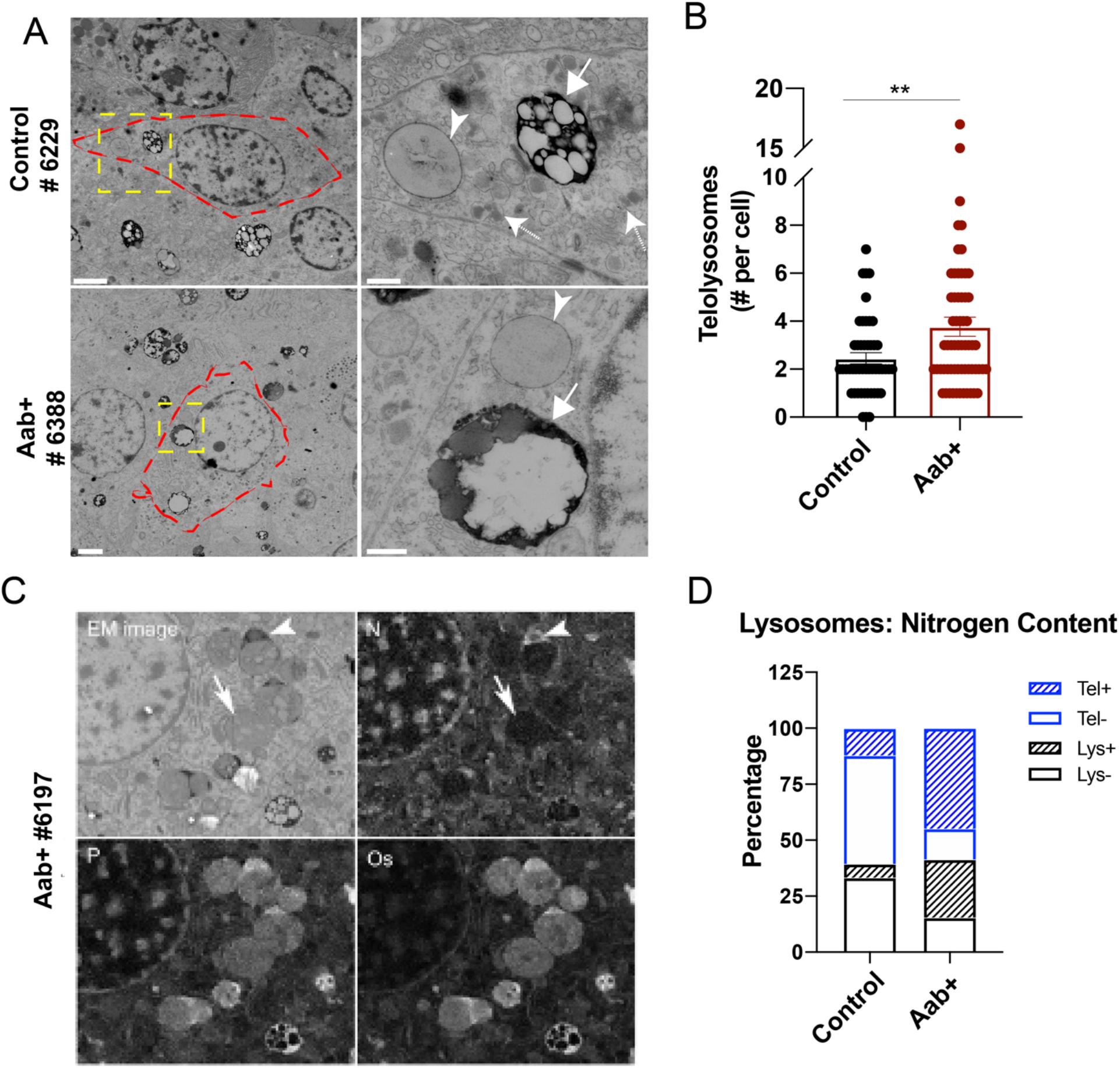
Increased number of telolysosomes in beta cells of autoantibody positive donors. **(A)** Representative images showing electron microscopic images of beta cells of nondiabetic, and autoantibody positive (Aab+) organ donors. Left panel: Scale bars, 2μm. Right panel represents zoomed in regions: Scale bars, 500nm. Arrow heads represent secondary lysosomes, arrows with solid line represent telolysosomes, and arrows with dashed lines represent insulin granules. **(B)** Quantification of telolysosomes. Each circle denotes an islet. **(C)** Representative images showing EDX analysis of elements-Nitrogen (N), Phosphorus (P), and Osmium (Os). Arrowheads represent telolysosomes with nitrogen ring in the rim, arrows represent telolysosomes without nitrogen rim **(D)** Quantification of percentage occupied by secondary lysosomes without nitrogen rim (Lys-), secondary lysosomes with nitrogen rim (Lys+), telolysosomes without nitrogen rim (Tel-), and telolysosomes with nitrogen rim (Tel+). *p<0.05; **p<0.01(unpaired student t-test).

### Elemental analysis of lysosomes and telolysosomes

We went on to perform elemental analysis to identify phosphorous, osmium, and nitrogen in beta cell lysosomes and telolysosomes from nondiabetic and autoantibody positive donors (**Fig. 5C**). We observed phosphorus colocalizing with osmium used in sample preparation to fix (membrane) lipids, suggesting presence of phospholipids in all samples. We also observed an increased percentage of lysosomes/telolysosomes with nitrogen accumulation associated with the phospholipid rich lysosome core in the beta cells of autoantibody positive donors (**Fig. 5C-D;** 70% of lysosomes + telolysosomes vs 18% in nondiabetic donors). Zinc and Sulphur, being components of insulin, were below detection limit in these lysosomes. The origin of this nitrogen rim remains unclear.

## Discussion

Type 1 diabetes pathogenesis is classically viewed with an immune-centric focus, due to the critical role for autoimmunity in beta cell apoptosis. However, recent evidence suggests early precipitating events in type 1 diabetes pathogenesis may originate within the islet itself, leading to targeting of the beta cell for destruction by the immune system [1,13]. Autophagy has been identified as an important player in beta cell survival and growth, contributing to cellular homeostasis and stress response [30]. While the role of autophagy in the context of type 2 diabetes has been studied extensively [31, 32], its role in type 1 diabetes is relatively unexplored. Here we show that clearance of beta cell autophagosomes is impaired in the context of human type 1 diabetes and in the NOD mouse model of diabetes, and that beta cell crinophagy is impaired in human type 1 diabetes. Therefore, we propose that impaired autophagy and/or crinophagy plays a role in type 1 diabetes disease pathogenesis.

Prior studies have hinted at defective autophagy in the context of type 1 diabetes. For example, a type 1 diabetes susceptibility gene, *CLEC16A,* was shown to stimulate rodent beta cell insulin secretion through the stimulation of a selective form of autophagy known as mitophagy, whereas humans with the susceptibility allele exhibited both reduced *CLEC16A* expression and increased HbA1c [33]. Another type 1 diabetes susceptibility gene, Cathepsin H *(CTSH)* [34, 35], is a lysosomal cysteine protease that plays a crucial role in lysosomal protein degradation. Importantly, the *CTSH* susceptibility allele is associated with faster disease progression in newly diagnosed diabetes and reduced β-cell function in healthy humans [36].

Our observation of reduced colocalization of LC3 and Lamp1 in both human type 1 diabetes and NOD mice suggests that the later stages of the autophagy cascade, where the autophagosome fuses with the lysosome, are likely impaired. This observation thus raises several questions: 1) Are there defects in vesicle fusion machinery leading to decreased autophagy in type 1 diabetes? 2) Is there a defect in transport of autophagosomes to lysosomes along microtubules? or 3) Are the lysosomes themselves mis-localized and not readily available for fusion? Lastly, 4) Is there a defect in lysosome function, perhaps due to localization or acidity [36, 37]?

Of the possibilities listed above, our data hints at the possibility of lysosome defects associated with type 1 diabetes pathogenesis. Lysosomes are major nutrient sensing organelles that not only function as a degradation system, but also play a major role in stress adaptation [38]. Defects in lysosomal function and/or acidity have been implicated in aging and in the pathogenesis of various diseases such as Parkinson’s, Huntington’s, Alzheimer’s, Gaucher, Nieman-Pick, mucolipidosis, Batten’s, and lipofuscinosis [39, 40]. The lysosomal protease, *CTSH*, provides a direct link between lysosome function and type 1 diabetes susceptibility, where the *CTSH* susceptibility allele is associated with impaired beta cell function in humans [36]. In support of the hypothesis that lysosome dysfunction may be involved in diabetes pathogenesis, genetic knockout of a key isoform of the vacuolar type H^+^ ATPase that is present on lysosomes led to impaired insulin secretion in mice [41]. Together, these studies suggest a crucial role for functional lysosomes in the regulation of beta cell homeostasis and function.

Electron Microscopy (EM) has long been a gold standard in autophagy studies. We analyzed beta cells from nPOD donor pancreata, and observed a significant increase in the number of telolysosomes, or lipofuscin bodies, in the beta cells of autoantibody positive donors. These represent lysosomes that have accumulated oxidized and highly crosslinked proteins, lipids and sugars that are undigestible. Our data suggest that in addition to their characteristic visual features that differentiate telolysosomes from normal secondary lysosomes, many can be further differentiated by the presence of a peripheral nitrogen accumulation associated with a phospholipid rich core. Of note, despite not being able to digest the lysosome contents, lipofuscin bodies constantly receive additional lysosomal enzymes from the cell in an attempt to digest the materials, ultimately further contributing to a relative deficiency of lysosomal enzymes in the cell [42]. Our observation of increased telolysosomes in the beta cells of autoantibody positive individuals, taken together with our observation of a trend towards increased autophagosomes in beta cells of autoantibody positive donors, hints that impairment in autophagy could occur before the development of clinical hyperglycemia in autoantibody positive individuals perhaps due to defective lysosomes. Collectively, these observations support a critically important role for beta cell lysosomes in the pathogenesis of type 1 diabetes.

In addition to impaired macroautophagy, we also observed reduced crinophagy in the beta cells of individuals with T1D. It was recently demonstrated that crinophagic bodies may contain short peptides with potentially immunogenic epitopes, suggesting that altered crinophagy could lead to the presentation of altered peptides that recruit T-cells to the beta cell [25]. Therefore, our data would suggest that although there is not globally decreased crinophagy in early type 1 diabetes pathogenesis (i.e., autoantibody positive donor beta cells), the processing of proteins within the crinophagic bodies is perhaps already altered, appearing as telolysosomes with undigested material by EM analysis.

Several limitations remain that must be addressed in future studies. For example, although our data is suggestive, in our current study we don’t fully answer the question of whether impaired autophagy or crinophagy is a cause or consequence of hyperglycemia. Furthermore, autophagy and crinophagy are dynamic processes, and our analyses represent only a static snapshot in time. It is clear that exogenous factors may dynamically change autophagic flux associated with lysosome dysfunction in the beta cell, as is the case for our data and evidence in the literature of the effects elicited by proinflammatory cytokines [43]. Therefore, while our data suggest globally impaired autophagy and crinophagy in human type 1 diabetes, perhaps linked to defective lysosome function, they are only suggestive and do not tell us if the kinetics are perturbed during diabetes pathogenesis. Since the data obtained from our NOD mice are concordant with our human data, NOD mice can potentially be used as a model to study type 1 diabetes associated autophagy impairment. Future studies that can incorporate analysis of dynamics in the intact mouse pancreas, such as novel non-invasive imaging approaches [44], or intravital microscopy approaches [45] to selectively study beta cell mass, function, and signaling at the single islet and subcellular level will be crucial to fully understanding autophagy defects in the context of type 1 diabetes pathogenesis.

## Conclusion

In conclusion, we provide evidence that beta cell autophagy and crinophagy are reduced in human type 1 diabetes. These results have potential clinical implications for type 1 diabetes prevention, as autophagy stimulators are available and currently in clinical trials (NCT03037437, NCT03309007). We anticipate that further studies of autophagy and crinophagy in the context of type 1 diabetes will yield additional insight into selective therapeutic targets that will add to this list in the future.

## Acknowledgements

We thank Dr. Sarah A. Tersey and Dr. Emily K. Sims for sharing of resources. We would also like to thank Dr. Raghu Mirmira and Dr. Manami Hara for their helpful discussions of this data. This research was performed with the support of the Network for Pancreatic Organ donors with Diabetes (nPOD; RRID:SCR_014641), a collaborative type 1 diabetes research project sponsored by JDRF (nPOD: 5-SRA-2018-557-Q-R) and The Leona M. & Harry B. Helmsley Charitable Trust (Grant#2018PG-T1D053). Organ Procurement Organizations (OPO) partnering with nPOD to provide research resources are listed at http://www.jdrfnpod.org//for-partners/npod-partners/. This work was supported by the Histology Core of the Indiana Center for Musculoskeletal Health at IU School of Medicine and the Bone and Body Composition Core of the Indiana Clinical Translational Sciences Institute (CTSI).

## Contributions

CM was responsible for acquisition, analysis and interpretation of data, and drafting the manuscript. AMC was responsible for analysis of data and revision of manuscript. MRM, JC, JK, PDB, and BNG were responsible for acquisition of data and revision of manuscript. AKL was responsible for conception and design, interpretation of data and drafting of manuscript.

## Funding

This research was performed using resources and/or funding provided by National Institutes of Health grants R03 DK115990 (to AKL), Human Islet Research Network UC4 DK104162 (to AKL; RRID:SCR_014393), along with startup funds from Indiana University School of Medicine and the Herman B Wells Center for Pediatric Research (to AKL).

## Conflict of interest

The authors declare no conflicts of interest.

## Abbreviations

nPOD: Network for Pancreatic Organ Donors with Diabetes
ROS: Reactive Oxygen Species
DSHB: Developmental Studies Hybridoma Bank
NOR: Nonobese Resistant
NOD: Nonobese Diabetic
CQ: Chloroquine
ROI: Region of Interest
EM: Electron Microscopy
EDX: Energy Dispersive X-ray Analysis

## Muralidharan et al. Supplemental Figures

**ESM Figure 1.**
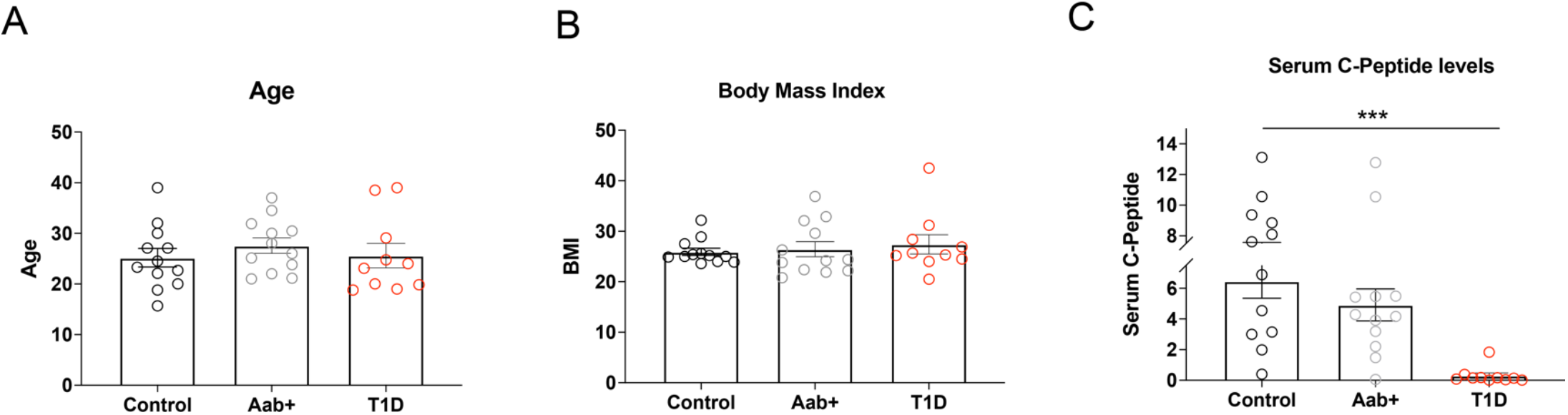
Characteristics of pancreatic tissue sections of donors used in the immunofluorescence staining experiments of human pancreatic tissues. **(A)** Age of the donors. **(B)** Body mass index of the donors. **(C)** Serum C-peptide levels of the donors.

**ESM Figure 2.**
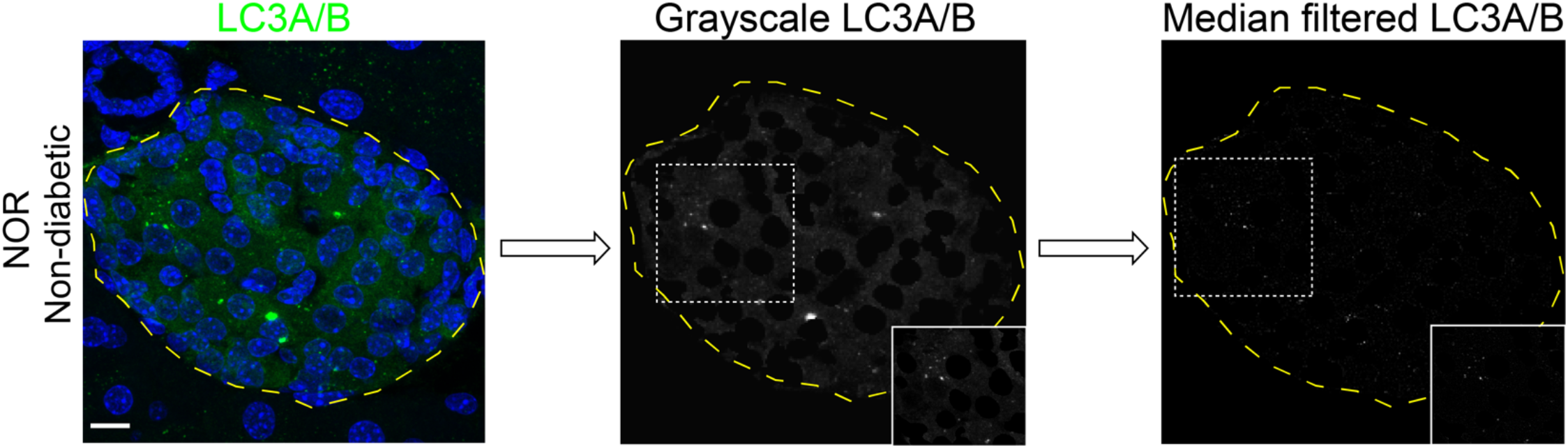
Example of identification of LC3 puncta after applying median filtering using CellProfiler. Scale bar, 10μm.

**ESM Figure 3.**
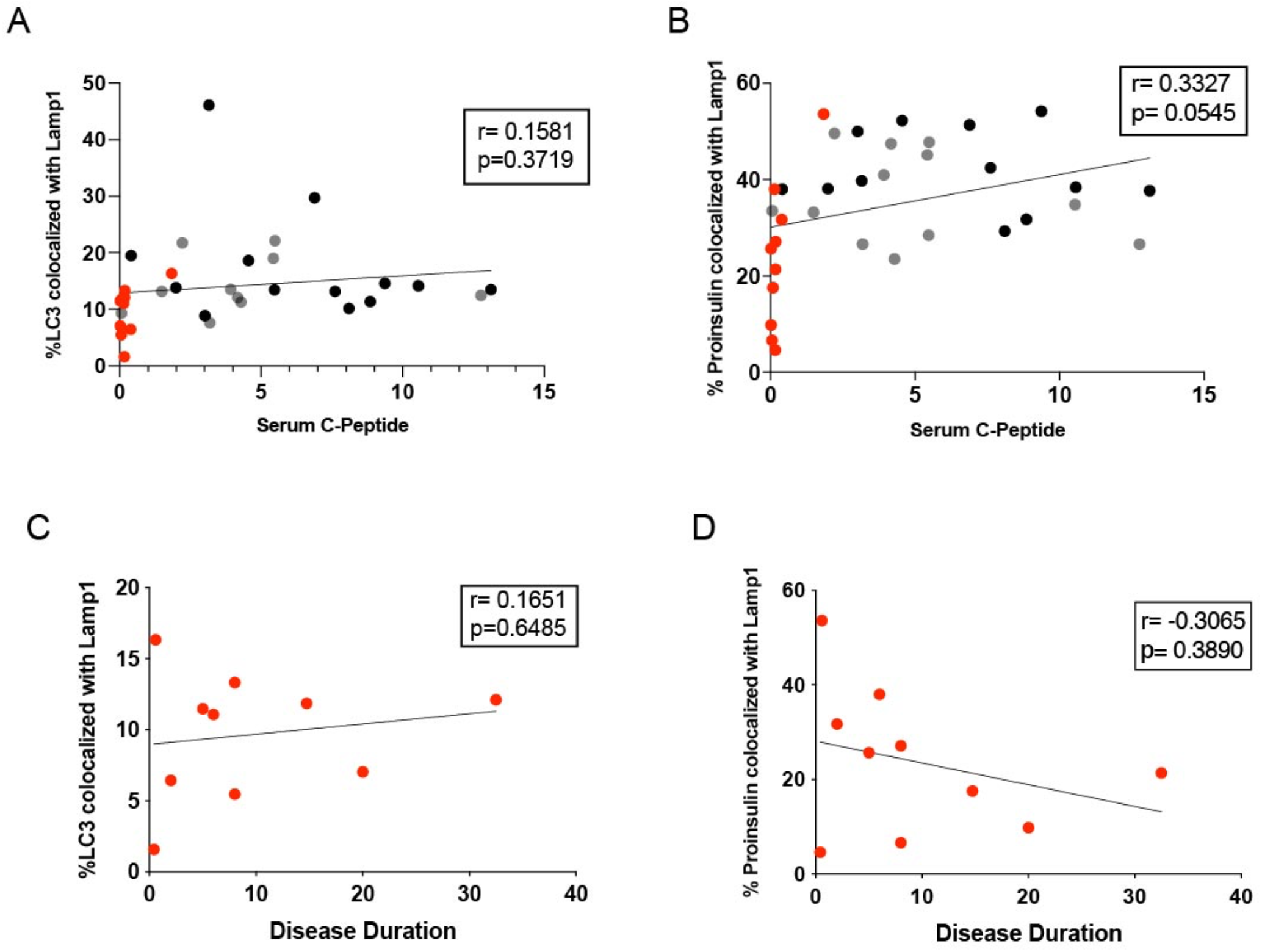
Correlation graphs of autophagy and crinophagy with serum C-peptide levels, and disease duration.

## Notes

### Competing Interest Statement

The authors have declared no competing interest.

